# Oxytocin and Emotional Tear in Dogs

**DOI:** 10.1101/2022.03.09.483532

**Authors:** K. Murata, M. Nagasawa, T. Onaka, S. Kanemaki, K. Nakamura, K. Tsubota, K. Mogi, T. Kikusui

## Abstract

In humans, tear volume increases in emotional arousal. No studies have investigated the relationship of emotional arousal and tear volume in animals. We measured tear volume in dogs before and after reunion with their owners. Tear volume increased significantly during owner, but not familiar non-owner, reunions but not during reunions with familiar non-owners. When oxytocin instillation was applied to dogs, the tear volume increased, suggesting that oxytocin can mediate the emotion-induced tear secretion in dogs. When the photos of dog’s face in which artificial tear was applied to the dog’s eyes, the positive impression of these photo increased. These suggest that emotional tear can facilitate human-dog emotional connection.

## Introduction

Dogs have been reported to have notable human-like social-cognitive skills, which are thought to result from convergent evolution with humans^1^, and they can use the eye to communicate with humans. For example, dog have evolved the eye muscle responsible for raising the inner eyebrow for triggering nurturing behavior in humans^2^. One of the most important skills is that dogs can use eye contact as a cue during communication with humans. Eye contact also played a pivotal role in attachment behavior in dogs. A dog’s gaze initiates interactions with its owner, and stimulates oxytocin (OXT) secretion in owners^3^, which is a key hormone to form a biological bond.

Humans often exhibit increased lacrimation, which appears as tears, in situations involving in pain^4^. Interestingly, humans also have emotional tears that are secreted in response to emotional arousal, both in positive and negative^6^. Tears also have a role in nonverbal visible communication, and are used when infants want to communicate negative feelings, such as hunger, pain, and discomfort, to their parents, and positive feelings, such as reunion with parents or friends^6^.

When dogs are reunited with their owners after a separation, they exhibit highly affiliative behavior, including gazing at their owners, wagging their tails, jumping up, and licking their owner’s faces, with changes in their autonomic functions^7^. Oxytocin levels rise in dogs during reunion with humans^8^ and then activate the parasympathetic nervous system^10^.

## Methods and Results

In this study, we hypothesized that dogs secrete emotional tear after reunion with the owner, and the tear secretion is controlled by oxytocin. We also examined whether the tears in dog’s eyes can facilitate human caregiving behavior toward dogs. At first, the tear volumes from before and after dogs were separated from their owners at their home or daily location were measured. A mixed model analysis revealed that tear volume was significantly increased during the reunions (Fig. 1A). Secondary, we examined whether the changes in tear volume were specific to reunions with the dog’s owner or not. After the separation from the owner in the dog’s day care centers, dogs showed more tear volume during reunions with the owner as compared to with the familiar non-owner (Fig. 1B). To reveal the OXT effects on tear secretion in dog, OXT solution was applied to the ocular surface and measured tear volume before and after the application. It was found that tear volume was significantly increased after OXT, but not control peptide, administration (Fig. 1C). These results showed that tear volume increased when the dogs reunited with their owner and this response was specific to the owner. In addition, because OXT is considered an important hormone in the establishment of biological bonding between owners and their dogs and secreted both in the owner and dog with interaction, our findings of increased tear volume due to the administration of OXT suggested that OXT stimulate the secretion of tear. In mice model, the function of OXT on the tear secretion is elucidated; OXT acts on the OXT receptor expressing on the myoepithelial cells in the lacrimal gland and oxytocin stimulates tear secretion^10^.

**Figure 1.**
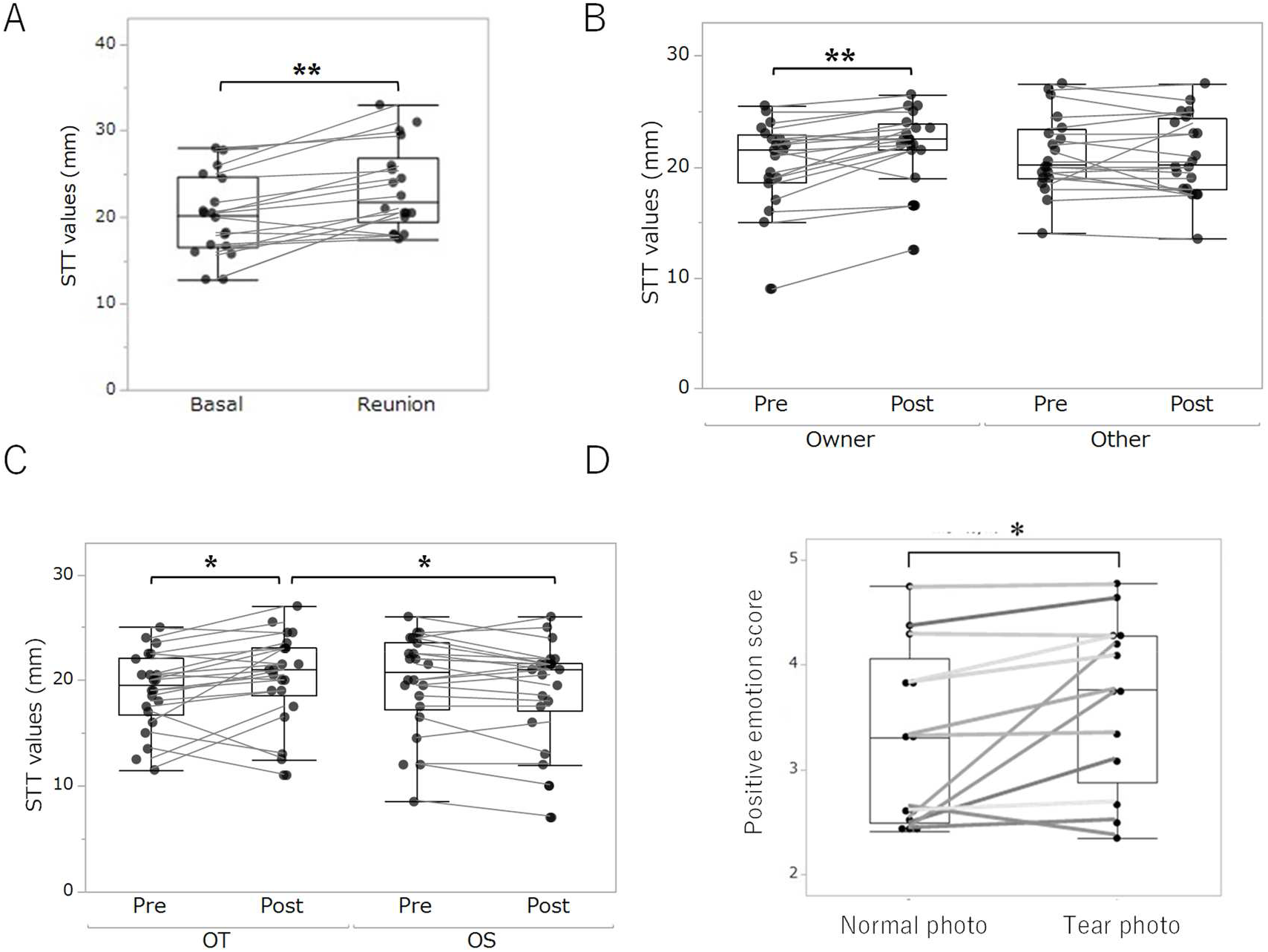
(A) Tear volumes was measured by Schirmer Tear test (SST), expressed the length of wetted part of the test paper. The baseline was measured when the dogs were stayed with owner in their house. Tears during the reunion was measured in dogs which was separated from the owner for more than 5 hours, and within the first 5 min of reunion. The tear volume during the reunion was larger than that of baseline (p<0.01, F(1,16)=8.54, Linear mix model). (B) Tear volume during the reunion with the owner and with the familiar non-owner was measured in dog’s day-care center. Pre volume was measured before the reunion, and Post was measured during the first 5 min of reunion of the dog which was separated from the owner for more than 5 hours. Tear volume during the owner reunion (Post), but not non-owner, was larger as compared with Pre (p<0.05, F(1,54)=4.35, Linear mix model). (C) Oxytocin (OT) stimulation for tear secretion. OT and control peptide (composed of the same amino acid components as OT, but the sequence was scrambled; OS) were dissolved in sterile saline (20 IU/120 μL) and dropped into the ocular surface of the both sides. The STT was conducted 5min after administration. Of the 22 animals, 11 received OT first, and the remaining 11 received OS first, to counterbalance the order of eye administration. The tear volume after OT administration was larger than that of OS and than Pre (p<0.05, F(1,60)=6.57, Linear mix model). (D) The impression scale of human participants after watching dog face photos. A pair of photos, one was normal face photo and the other was face photo in which sterile saline was applied to the eyes of the same dogs, was prepared for each dog. 10 photos from 5 dogs were randomly presented on the screen of PC, and the participants scored on a 5-point scale from positive (want to touch, and give some cares) to negative (fearful and avoid) for each photo. The photos of face with tears were judged to be more positive than the photos of the normal face (p<0.01, Wilcoxon signed-rank test).

Finally, cognitive function of tear secretion in dogs were evaluated. It is hypothesized that the emotional tear in dogs can facilitate human caregiving to dogs, as in the human children. The photos of face with tears were judged to be more positive than the photos of the normal face (Fig. 1D).

## Discussion

In this study, we demonstrated that dog secret emotional tear when reunite the owner, and this tear secretion is suggested to be mediated by oxytocin, if the dogs were treated with oxytocin, the tear fluid volume increase. This is the first report demonstrating positive emotion stimulate tear secretion in non-human animals, as well as the OXT function in tear secretion.

In humans, social and emotional status are associated with tear secretion. When the prefrontal brain regions that govern emotions are activated, the neural information are transmitted to the superior salivatory nucleus, which causes parasympathetic hyperactivity^13^. This subsequently results in the secretion of large volumes of tears from the lacrimal glands that then overflow from the eyes. Recently we discovered the OXT can facilitated the synthesis and release of tears in the mouse brain; OXT neurons located in the paraventricular hypothalamus send the fibers to the superior salivary nucleus and increase the tear secretion. Therefore, OXT increase during the reunion with the owner can act on both central and peripheral organs to stimulate tear secretion^14^.

The social functions of OT-stimulating tear are unsolved at this moment. Tears are used as chemosignals for communication in mammals. For example, male mouse tear contains ESP1, which stimulates female sexual behavior and male aggression^15^. Dogs may use the tear as commutation signals, for example, sniffing and licking near the eyelid is commonly observed in dogs’ greeting^12^. Even in humans, tear fluid contained some chemosignals and tears is used as socio-sexual signals^16^. Therefore, one important function of tear fluid is chemo-signaling in mammals. In humans, social and emotional status can be transmitted by tear^14^. Infants use tears to communicate feelings, especially negative ones such as hunger, pain, and discomfort, to their parents^15^. Therefore, tears can act as an attachment signals from the sender, and the receivers show caregiving behavior. In this study, we found that tearing dog face can facilitate human caring emotion to dogs, which is similar function observed in human children. We also discovered that emotional tear can be also observed in mice, maternal reunion, nociceptive stimulation, and aversive memory retrieval are linked to tearing in mice^14^. The social function of tearing in animals needs to be elucidated in future.

Unlike any other animal, dogs have evolved or domesticated through communication with humans. Compared to primates, such as chimpanzees, the closest living relative of humans, dogs have a superior ability to understand social signals, such as eye gaze and gestures of humans^1^. This is thought to be due to dogs having superior social-cognitive abilities that enable communication with humans as a result of sharing a niche with them. The interactions between owners and dogs are initiated by the gaze of dogs, and urinary OXT levels increase in owners after interactions with their dogs^3,17^. These findings suggest that eye gaze plays a significant role in the formation of bonds and communication between humans and dogs. The tears of dogs, which have evolved high-level communication abilities with humans through the use of eye to eye communication, might also function to elicit protective behavior or nurturing behavior from their owners, resulting in the deepening of mutual relationships, which led to interspecies bonding.

## Acknowledgments

We thank Ms Naoko Tsuchihashi of Azabu University and veterinary medical staffs in Mominoki Veterinary Clinic and Inclover-world in Hyogo, Japan, for help in behavioral experiments.

## Author Contributions

K.Murata, M.N., K.Mogi and T.K. designed the research. K.Murata and M.N. conducted behavioral experiments. T.O. and K.Mogi. conducted hormonal experiment. K.K. and K.Mogi developed measuring tear. T.K., K.M. and M.N. conducted statistical analysis. K.Murata and M.N. drafted the manuscript, and M.N., K.Mogi, S.N, K.Tsubota and T.K. edited it. All authors read and approved the manuscript.

## Data and Materials Availability

Data can be shared by the corresponding author upon the request.

